# Impact of substrate-template stability, temperature, phosphate location, and nick-site base pairs on non-enzymatic DNA ligation: Defining parameters for optimization of ligation rates and yields with carbodiimide activation

**DOI:** 10.1101/821017

**Authors:** Chiamaka Obianyor, Gary Newnam, Bryce Clifton, Martha A. Grover, Nicholas V. Hud

## Abstract

Non-enzymatic, chemical ligation is an important tool for the generation of synthetic DNA structures, which are used for a wide range of applications. Surprisingly, reported chemical ligation yields range from 30% to 95% for the same chemical activating agent and comparable DNA structures. We report a systematic study of DNA ligation using a well-defined bimolecular test system and water-soluble carbodiimide (EDC) as a phosphate-activating agent. Our results reveal interplay between template-substrate stability and the rates of the chemical steps of ligation, which can cause yields to increase or decrease with increasing temperature. Phosphate location at the nick site also exhibits a strong influence on ligation rates and yields, with a 3’ phosphate providing yields near 100% after 24 hours for particularly favourable reaction conditions, while comparable reactions with the phosphate on the 5’ position of the nick site only reach 40% ligation even after 48 hours. Ligation rates are also shown to be sensitive to the identity of base pairs flanking a nick site, with some varying by more than three-fold. Finally, DNA substrate modification by EDC can, in some cases, make long reaction times and repeated addition of EDC an ineffective strategy for increasing ligation yields.

## INTRODUCTION

Naturally derived ligase enzymes are widely used for joining DNA strands that are isolated from living organisms, prepared by the polymerase reaction, and synthetized using solid-state methods. T4 DNA ligase, for example, is used extensively as part of many procedures in recombinant DNA technology. Despite its incredible utility, enzymatic ligation is limited by specific reaction conditions that are optimal for a particular ligase enzyme. Furthermore, naturally derived ligases are highly selective for Watson-Crick DNA duplexes and are therefore not typically suitable for the ligation of DNA duplexes containing non-Watson-Crick base pairs, non-duplex structures, and enzyme-inaccessible regions, all of which can appear in structures designed for DNA nanotechnology (1,2). For such applications non-enzymatic chemical ligation offers potential advantages over enzymatic ligation, as well as lower cost, which could be important for large-scale reactions. Additionally, DNA is being used for the exploration of minimal, enzyme-free self-replicating systems, which would benefit greatly from the development of efficient chemical ligation protocols that can operate under conditions needed to cycle DNA templates and substrates through multiple rounds of replication (3–5).

To our knowledge, there is no consensus on which chemical activating agents are most effective for non-enzymatic DNA ligation, or whether one particular buffer, catalyst, or temperature is optimal for most ligation reactions. Additionally, reported chemical ligation yields range from as low as 30% to as high as 95% for the same chemical activating agent and for seemingly comparable DNA structures (6). Cyanogen bromide, which has been used for ligation in template-directed DNA synthesis (6–8), is one of the simplest and longest recognized phosphate-activating agents. However, the rapid kinetics of cyanogen bromide hydrolysis and incomplete ligations achieved with even high concentrations of this highly toxic reagent suggest that it would be difficult to further develop this reagent as an optimal activating agent. Cyanoimidazole (a milder activating agent) allows for longer reaction times than cyanogen bromide, but has also shown limitations for achieving quantitative ligation (9). Previous reports have shown that DNA ligation using water-soluble carbodiimides can approach 95 %, after 6 days (6,10,11), but most reported yields fall far short of quantitative ligation and are severely impacted by side reactions (4). The optimization of RNA chemical ligation using water-soluble carbodiimides has been explored, but the yields of RNA chemical ligation also remain low (12,13). Thus, there remains a need for a rubric by which nucleic acid chemical ligation protocols can be designed to achieve optimum product formation under a wide variety of conditions. Towards this end, we explored the factors that govern ligation rates and yields with soluble carbodiimides as activating agents (13,14).

Here we report the investigation of several experimental parameters that could limit DNA chemical ligation rates and overall yields; including the stability of the ligation complex, nick sequence specificity, impact of temperature on the chemical steps of ligation, and phosphate position at the ligation site (the duplex nick). Using a two-oligonucleotide ligation system with variations at the ligation site, we studied the interplay between hybridization stability, temperature, and ligation kinetics. We show that the position of the phosphate has a major impact on chemical ligation rates, with a 3’ phosphate being superior to a 5’ phosphate at the backbone nick. We also observed that the specific bases flanking a nick site can significantly impact the rate of ligation, particularly at lower reaction temperatures. Altogether, the results presented here should be useful for improving the synthesis of a wide range of synthetic DNA structures, which will hopefully facilitate the development of covalent structures for DNA nanotechnology (15), molecular sensing probes (16,17), and more.

## MATERIAL AND METHODS

### Oligonucleotide preparation

All oligonucleotides were ordered from Integrated DNA Technologies (IDT) and resuspended in 18.2 MΩ/cm water (Barnstead NanopureTM). The 3’p hairpin oligonucleotides were further purified using denaturing polyacrylamide gel electrophoresis. The 5’p hairpin oligonucleotides were prepared using T4 Polynucleotide Kinase (New England Biolabs).

### Ligation experiments

For a standard reaction, the fluorescently-labeled hairpin oligonucleotide was present at 1.3 μM along with the substrate oligonucleotides (at concentrations ranging from 0.5x to 50x hairpin concentration), in a buffer containing 5 mM MnCl_2_, 100 mM MES, pH 6.0, and 250 mM (1-Ethyl-3-(3-dimethylaminopropyl) carbodiimide) EDC (Sigma-Aldrich). Before addition of EDC, the oligonucleotides and buffer mixture were heated to 80 °C for 2 min, then quickly cooled to room temperature. Five minutes after cooling, EDC was added to the reaction to initiate the reaction, and the reaction tube was immediately moved to the specified temperature. After the indicated reaction time each reaction was quenched by the addition of an equal volume of 2x loading buffer and dye (95% formamide, 0.025% bromophenol blue, 0.025% xylene cyanol, 5 mM EDTA pH 8.0). Each sample was then stored at −80 °C until analyzed by denaturing polyacrylamide gel electrophoresis.

Polyacrylamide gels (Fisher BioReagents^TM^ acrylamide/bis-Acrylamide 29:1, 40% solution) were 20% denaturing gels (8 M urea) run in 1x TBE (Tris, Boric Acid, and EDTA pH 8.0) buffer, 5 cm wide X 15 cm long. Gels were pre-run at 14 W and 300-400 V for at least 30 min prior to loading. Samples were run at the same conditions for 1 h. Imaging was done using a Typhoon Trio+ laser scanner (GE Healthcare) at a resolution of 50 μm and with a photomultiplier setting between 300-500. ‘FAM filter’ images were acquired using the ‘FAM channel’, which refers to 488 nm excitation and a 526 nm emission filter. Densitometry analysis was performed using utilities within the ImageJ software package (NIH).

### Densitometry analysis of ligated products

To obtain % ligated products, the integrated intensity (densitometry measurement) of each gel band corresponding to a ligation product was divided by the sum of the integrated intensity of the ligation product and unreacted hairpin oligonucleotide in each lane. These % products are used to describe all ligation results.

## RESULTS AND DISCUSSION

### Design of a ligation test system

Our ligation test system is based on 28-nt hairpin-containing oligonucleotides that are labeled on their 3’ ends with 6-FAM (Fluorescein). These oligonucleotides, referred to throughout simply as “hairpins,” also contain a 9-nt single-stranded overhang that serves as a substrate binding site for template-directed ligation (Scheme 1A). Two sets of substrate oligonucleotides were chosen, a 9-mer set, which was fully complementary to the substrate binding site of the hairpin, and a 5-mer set, which was complementary to a subsection of the same substrate binding site. These substrate oligonucleotides were designed to explore non-enzymatic ligation within stable (or partially stable) pre-ligation oligonucleotides assemblies (i.e., hairpin-substrate assemblies). The substrate oligonucleotides are longer than those for which intramolecular cyclization is typically favored over ligation (i.e., dimers, trimers, and tetramers (13,18)), so competition between ligation and cyclization is minimized (particularly in the 9-mer system). The phosphate group at the nick site of the hairpin-substrate assembly was on the 3’ terminus of the substrate oligonucleotides for most investigations reported here because ligation rates and yields were found to be higher than if the phosphate at the nick site was on the 5’ terminus of the hairpin (see below).

For all reactions, the water-soluble condensing agent EDC was used for *in situ* activation of the oligonucleotide terminal phosphate group. EDC was added to pre-mixed solutions containing hairpin and substrate oligonucleotides, and the reaction tubes were immediately moved to one of three incubation temperatures (4, 25, or 37 °C). The hairpin oligonucleotides were present at 1.3 μM in all reactions unless otherwise noted. Concentrations of 9-mer and 5-mer substrates are provided in relative molar ratios to the hairpin (ranging from 0.5:1 to 50:1, as noted). Given that the half-life of EDC in solution is around 16 hr (4,14), we were particularly interested in identifying reaction conditions that would result in full ligation within 24 hr, or 48 hr, if complete ligation within 24 hr was not possible. Preliminary studies revealed that buffer conditions previously optimized for DNA ligation with cyanoimidazole activation (18) were also near optimal for ligation with EDC activation, and did provide near quantitative ligation for some ligation systems within the desired time frame. Thus, these reaction buffer conditions of 5 mM MnCl_2_, 100 mM MES buffer (pH 6.0), and 250 mM EDC were used throughout, unless otherwise noted.

Prior to covalent bond formation substrate oligonucleotides can exchange between solution and the substrate binding site of the hairpins (Scheme 1A), governed by an equilibrium constant that depends on several factors, including substrate length, hairpin/substrate concentrations, ionic strength, and temperature. Activation of the substrate by EDC is depicted in Scheme 1B for a free substrate oligonucleotide, but activation could also happen while the substrate oligonucleotide is bound to the hairpin (not depicted in Scheme 1). The final step of ligation, formation of the phosphodiester bond at the nick site (Scheme 1D), is considered irreversible. The last three reactions depicted in Scheme 1 (E-G) represent side reactions that can reduce ligations rates and yields, including hydrolysis of the EDC leaving group from an activated substrate (deactivation) (Scheme 1E). EDC-induced reactions can inhibit substrate and hairpin from participating in the ligation reaction, such as cyclization, in the case of the substrate, or base modification for either substrate or hairpin oligonucleotides that interfere with hairpin-substrate association (Scheme 1F and 1G). These modifications have previously been noted and investigated by others (4,13,19).

### Interplay between template-substrate stability, temperature, and ligation rate

Substrate oligonucleotides were selected to test the impact of the hairpin-substrate complex stability on ligation rates and yields. In particular, we sought a set of substrate oligonucleotides that would be fully bound to the hairpin at and below 25 °C, and at least partially associated at 37 °C, the highest temperature used for our ligation reactions. A set of shorter substrates was also desired that would be partially associated with the hairpin at the lowest reaction temperature, 4 °C, but primarily or even completely dissociated at 25 °C and 37 °C. Based on predicted melting temperature (T_m_) values, 9-mer and 5-mer substrates were selected.

**Scheme 1.**
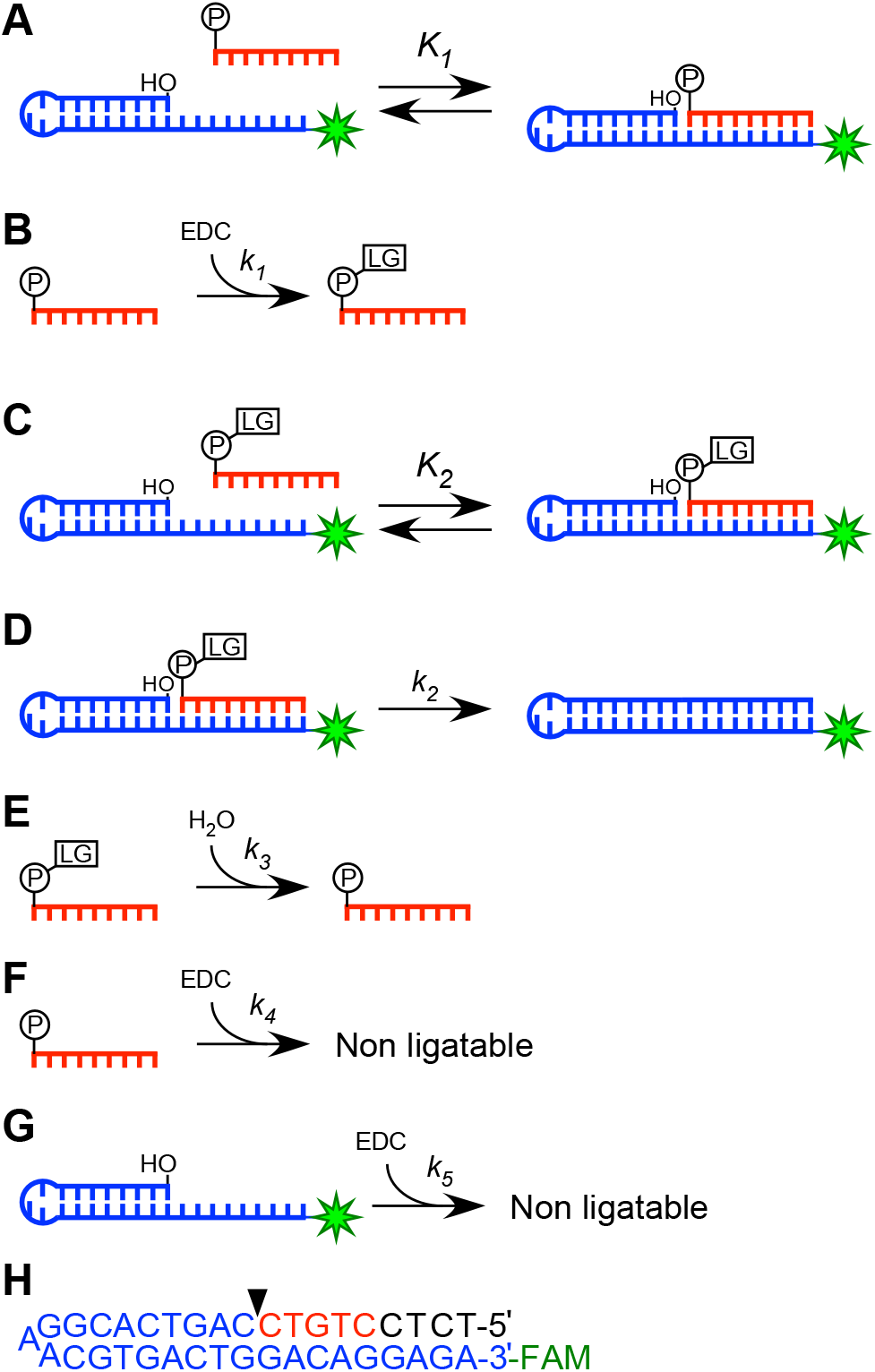
Illustration of the ligation test system with the equilibria and reactions that govern the overall rates and yields of ligation. A) Equilibrium of the hairpin (unactivated) substrate complex. B) Activation of the substrate oligonucleotide by EDC. This reaction is shown for a substrate oligonucleotide free in solution, but may also occur while the substrate is bound to the hairpin. C) Equilibrium of hairpin and activated substrate complex. D) Formation of the phosphodiester bond. E) Hydrolysis of activated substrate. F) EDC induced reaction that renders a substrate oligonucleotide non-ligatable (e.g., cyclization). G) EDC induced reaction that renders a hairpin template unable to participate in ligation (e.g., base modification that prevents substrate binding). H) Sequence and secondary structure of FAM-labeled hairpin (blue nucleotides); 5-mer substrate (red nucleotides), and 9-mer substrate (red and black nucleotides). Wedge indicates position of backbone nick at which the phosphate can be either on the 5’-terminus of the hairpin or on the 3’-terminus of the substrate oligonucleotide. Sequences shown are used in all ligation studies, except in the final experimental section where the impact of sequence variations is explored.

In Figure 1 we present results for the ligation of a particular 28-nt hairpin with its complementary 9-mer and 5-mer substrates as a function of time and temperature (hairpin and substrate sequences provided in Scheme 1H). For this set of reactions, the substrate oligonucleotides were present in a 10:1 relative molar concentration to the hairpin (i.e., 13 μM substrate, 1.3 μM hairpin) to ensure that the substrate oligonucleotide would not be a limiting reagent of the reaction, a point addressed in detail below.

**Figure 1.**
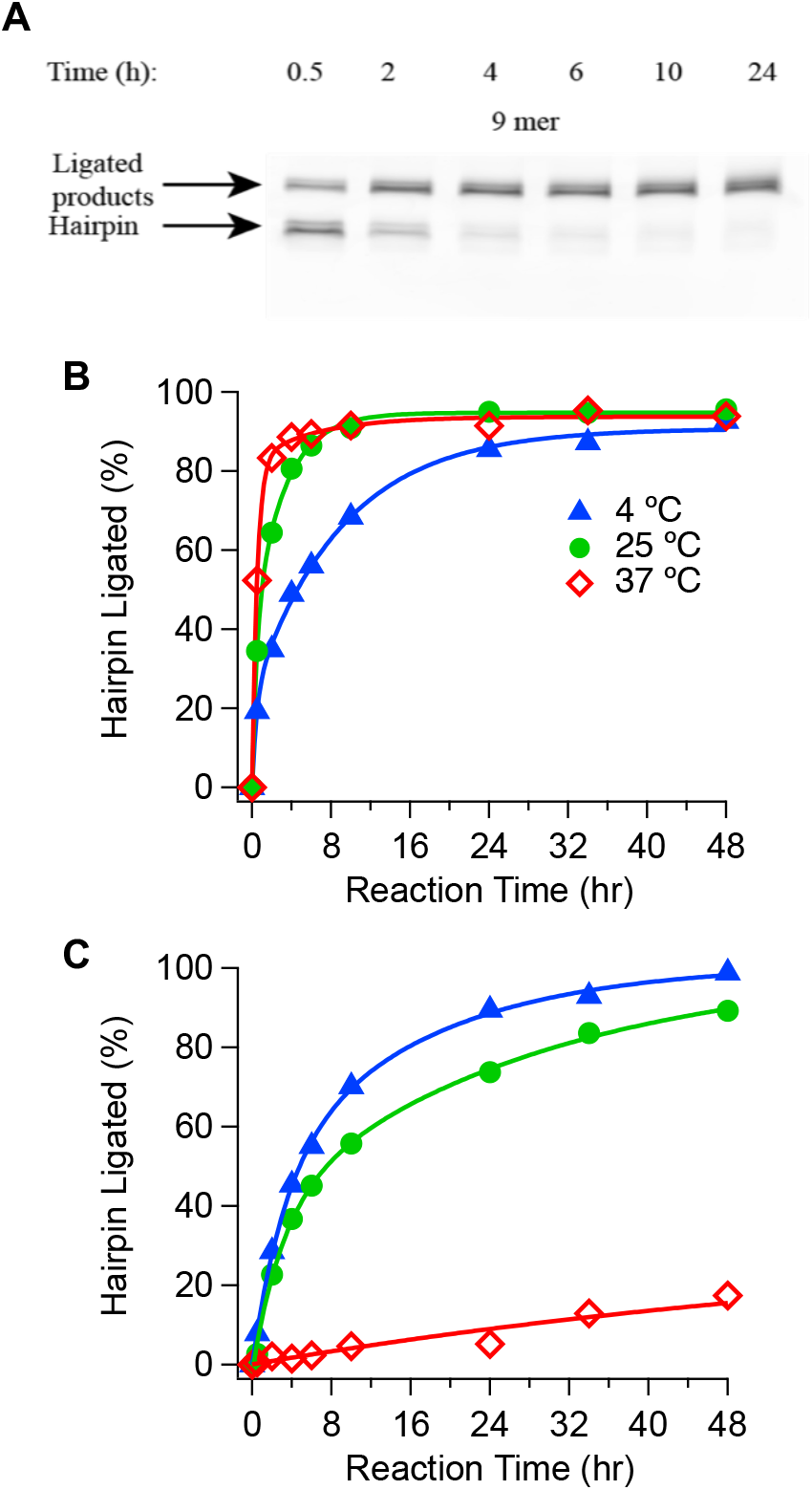
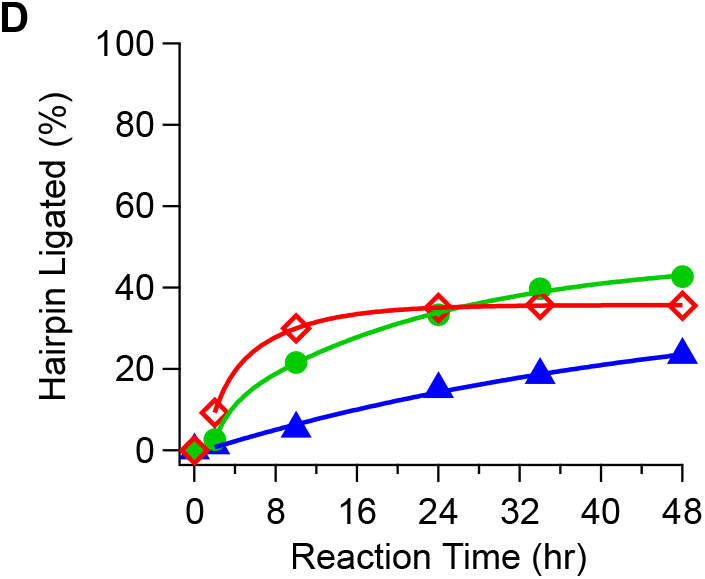
Kinetic measurements of chemical ligation at different temperatures. (**A**) Sample gel electrophoresis image of kinetic studies conducted at 25 °C for the 3’-phosphate 9-mer. Buffer conditions: 5 mM MnCl_2_, 100 mM MES, pH 6.0, and 250 mM EDC. (**B**) Kinetic data for hairpin ligation with the 3’-phosphate 9-mer substrate. (**C**) Kinetic data for hairpin ligation with the 3’-phosphate 5-mer substrate. (**D**) Kinetic data for 5’-phosphate hairpin ligation with the 9-mer substrate. 1.3 µM hairpin and 13 µM of substrate DNA oligonucleotides were used for all reactions shown. Data points were obtained by scanning of polyacrylamide gels, such as the example shown in Panel **A** (See Materials and Methods for additional information). The lines are double exponential fits to help guide the eye through the data points as well as to provide a visual comparison of relative initial reaction rates and the time at which each set of reactions reached 50% yield.

The ligation kinetics and yields observed for reactions carried out at 4 °C are quite similar for the 9-mer and 5-mer substrates. Both systems reached 50% ligation yield after reacting for 5 hr, around 80% yield at 24 hr, and over 90% yield at 48 hr (Figure 1B and C). Increasing the temperature of the 9-mer substrate reactions to 25 °C and 37 °C resulted in a substantial increase in ligation rates, with 50% ligation being reached within 1 hr, and maximum ligation yields of around 90% for both temperatures being achieved near the 10 hr time point. In contrast, increasing the temperature of the 5-mer substrate reactions to 25 °C caused a minor decrease in the ligation rate, and a small decrease in the yield measured at 48 hr. At 37 °C the ligation rate of the 5-mer substrate reaction was decreased so drastically that only 17% yield was measured at the 48 hr time point (Figure 1C).

The opposite response of the 9-mer and 5-mer substrate ligation reactions to increased temperature illustrates the interplay of two fundamental factors that govern ligation rates in our ligation test system and, presumably, in many published studies involving DNA chemical ligation. For the 9-mer substrate reactions the higher ligation rates at 25 °C and 37 °C can be attributed to an increase in the rate of the chemical step of ligation with temperature, which could include both the rate of substrate activation and the rate of hairpin-substrate phosphodiester bond formation (Scheme 1B and 1D). As mentioned above, the 9-mer substrate was chosen so that the pre-ligation hairpin-substrate assembly would be fully assembled up to at least 37 °C. Thermal denaturation studies confirmed that the T_m_ of the 9-mer substrate-hairpin complex is around 45 °C (Supplementary Figure S1). Thus, increasing the reaction temperature from 4°C to 37 °C did not result in an appreciable decrease in the fraction of hairpins with bound substrates in these reactions. Given that nearly 100% of the hairpins in these reactions have bound substrates, the ligation rates measured with 9-mer substrates at 25 ºC and 37 ºC may be near the maximum possible with EDC activation at these temperatures.

The rates of the chemical steps of the 5-mer substrate reactions are also expected to increase with temperature. Thus, the observed *decrease* in 5-mer substrate ligation rates at 25 °C and 37 °C is likely due to a decreased fraction of hairpins with bound 5-mer substrates at these higher reaction temperatures. The temperature dependent stability of the 5-mer substrate-hairpin assembly supports this possibility. Given that the initial rates for ligation of 5-mer and 9-mer systems at 4 °C are similar (approximated by the amount of product formed at 2 hr), it follows that the fraction of bound hairpin is significant for both substrates at that temperature. However, as the temperature is increased, the initial rate of the 5-mer ligation reaction decreases relative to the 9-mer ligation reaction by over a factor of 3 by 25 °C and by a factor of 40 by 37 °C, indicating that the relative fraction of hairpins with bound substrates drastically changes for these two substrates over the temperature range of 4 °C to 37 °C. This correlation is consistent with the predicted T_m_ of the 5-mer-hairpin and the 9-mer-hairpin complexes being around 6 °C and 45 °C, respectively.

To provide additional support for this possible explanation for the different response of the 9-mer and 5-mer ligation reactions to increasing temperature we used both thermal denaturation studies and duplex stability calculations to estimate the fraction of hairpin with bound substrates in these 10:1 substrate:hairpin reactions (Supplementary Table S1). Consistent with our ligation rate data, thermal denaturation studies indicate that 100% of the hairpin in the 9-mer ligation reactions at 4 °C and 25 °C substrate had bound substrate, and approximately 90% at 37 °C. In contrast, for the 5-mer around 60% of the hairpin is predicted to have had bound substrate at 4 °C, approximately 2% at 25 °C, and less than 0.1% at 37 ºC. These temperature-dependent substrate-hairpin complex stabilities illustrate the direct impact that changes in ligation complex stability can have on ligation reaction rates over a relatively small temperature range (e.g. 25 °C to 37 °C). It is important to note that these estimates for equilibrium constants are based on unactivated 9-mer and 5-mer substrates (Scheme 1A). EDC has been shown to stabilize pre-ligation DNA duplexes (4), and it is possible that the T_m_ values for the binding of the activated 9-mer and 5-mer substrates with the hairpin (Scheme 1C) will be different for EDC-activated substrates, a possibility that is given further support below.

As mentioned above in the description of our ligation test system, most reaction studies presented here involved 9-mer and 5-mer substrates phosphorylated on their 3’ termini. This placement of the phosphate at the site of the hairpin-substrate nick was chosen based on the observation that placement of the phosphate on the 5’ terminus of the hairpin resulted in substantially lower ligation rates and yields. As one illustration of this fact, Figure 1D shows data from reactions with the phosphate on the 5’ terminus of the hairpin under identical conditions to those presented in Figure 1A for reactions with the phosphate on the 3’ terminus of the 9-mer substrate. This change in phosphate position results in a substantial reduction in ligation rates at all temperatures tested, and in overall yield at the 48 hr time point. We expect that this substantial difference in ligation rates is due, at least in part, to the reactions with the phosphate on the 3’ side of the nick site benefitting from the ligation reaction nucleophile being a primary alcohol, whereas the reactions with the phosphate on the 5’ side of the nick site relying on a secondary alcohol as the reaction nucleophile. It is also possible that differences in reaction site geometries for the two phosphate positions could also contribute to the observed differences in reaction rates and yields. Additionally, placement of the phosphate on the hairpin could make the hairpin more susceptible to modifications by EDC that render the hairpin non-ligatable. These modifications would impact yields more directly in our experimental design than loss of ligatable substrate (which was initially at a 10:1 excess in these reactions), as modifications in a 3’ terminus phosphate substrate were found to severely impact yields for reactions with a 1.25:1 substrate:hairpin ratio (Figure S2).

### Impact of an organocatalyst and increased monovalent cation concentration on reaction rates

A recent study by Richert and co-workers demonstrated that 1-Ethylimidazole (1-Ethyl) greatly increased the ligation yields of RNA dinucleotides and trinucleotides in reactions with EDC as the phosphate activating agent (13). The same group had previously demonstrated that mononucleotides would undergo template-directed polymerization when activated by EDC only if 1-Ethyl or a similar organocatalyst was present in the reaction buffer (12). Based on these reports, we decided to investigate the impact of 1-Ethyl on our DNA ligation reactions. In Figure 2 we present the results from reactions that are identical to those shown in Figure 1, except in which 1-Ethyl was added to the ligation buffer along with EDC. Contrary to expectations, 1-Ethyl actually caused a reduction in the reaction rates for both the 9-mer and the 5-mer ligation reactions. The 9-mer substrate reactions carried out at 25 °C and 37 °C might be expected to reach yields comparable to reactions without 1-Ethyl, but we estimate these levels would require approximately three times as long (Figure 2A). This suppression of reaction rates was even more dramatic for the 5-mer-ligation system, with ligation rates and 48 hr yields falling by almost an order of magnitude (Figure 2B).

**Figure 2:**
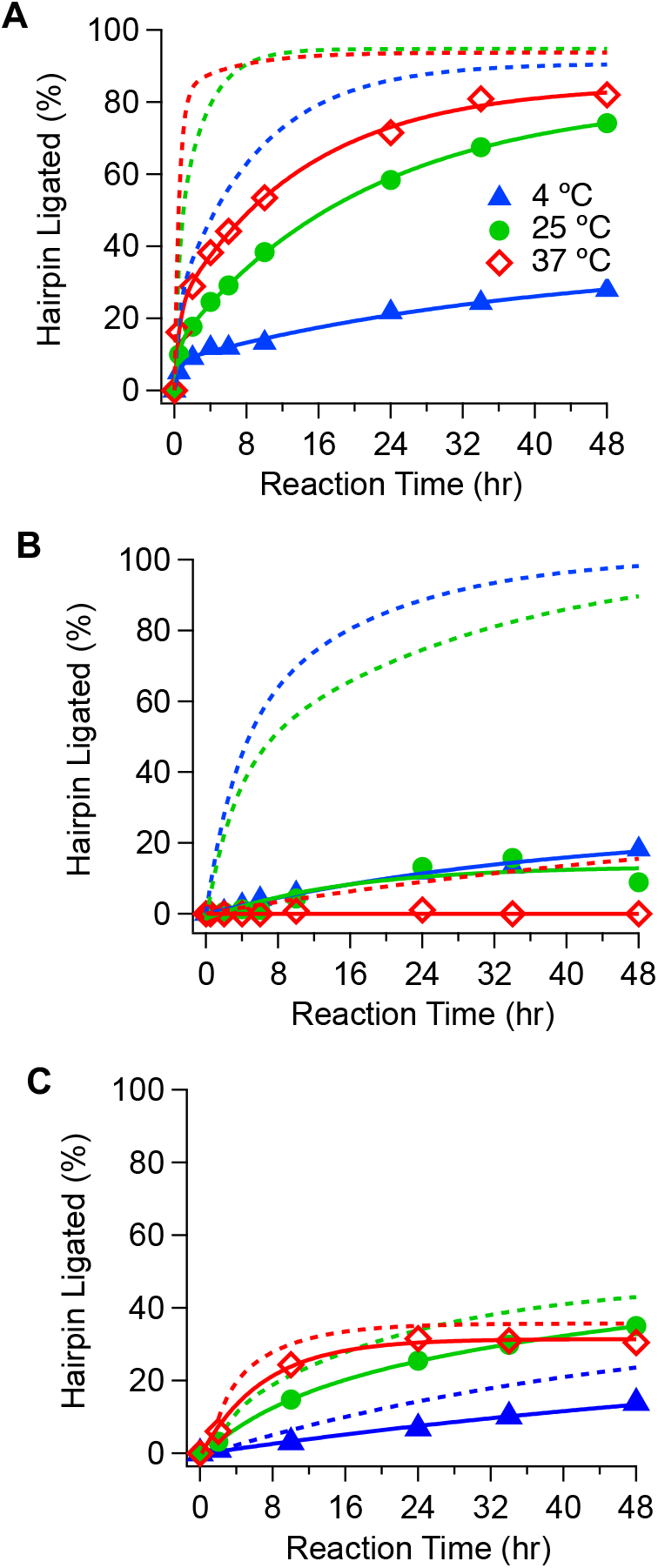
Effect of adding 1-Ethylimidazole to ligation reactions. (**A**) Kinetic data for hairpin ligation with the 3’-phosphate 9-mer substrate. (**B**) Kinetic data for hairpin ligation with the 3’-phosphate 5-mer substrate. (**C**) Kinetic data for 5’-phosphate hairpin ligation with the 9-mer substrate. Markers indicate experimental results for reactions with the addition of 1-Ethyl. Solid lines are double exponential fits of data, as described in Figure 1. Dashed lines are from the data of corresponding reactions shown in Figure 1. All data points were obtained as described in Figure 1 caption. Reaction buffer: 5 mM MnCl_2_, 100 mM MES, pH 6.0, 150 mM 1-Ethylimidazole (1-Ethyl), pH 6.0, and 250 mM EDC.

There are at least two important differences between our ligation reactions and those for which Richert and co-workers showed that 1-Ethyl catalyzes EDC-activated phosphate bond formation. First among these differences are that the current study is focused on the ligation of oligonucleotides with the phosphate on the 3’ terminus, whereas Richert and co-workers studied mononucleotide polymerization and ligation with the phosphate on the 5’ position. To test if phosphate position plays a role on the effectiveness of 1-Ethyl as a catalyst for phosphodiester bond formation we also ran a set of 9-mer ligation reactions similar to those of Figure 1D, but with the phosphate on the 5’ end of the hairpin as opposed to on the 3’ end of the 9-mer substrate. As shown in Figure 2C, the addition of 1-Ethyl to these reactions has a less dramatic impact, but the ligation rates observed up to 48 hr are still less than without 1-Ethyl added.

A second difference between our systems and those of Richert and co-workers is the use of DNA versus RNA, with the possibility that sugar structure influences the effectiveness of 1-Ethyl as a catalyst, a possibility that is not explored here. In any case, given the fact that 1-Ethyl did not enhance the ligation rates or yields of our DNA systems, we did not include 1-Ethyl in any other reaction reported in this study.

Reaction buffer ionic strength and ionic species are other possible modulators of ligation rates and yields. Although the buffer used for the ligation reactions reported here (5 mM MnCl_2_ and 100 mM MES, pH 6) was found to be optimal by the criteria of an earlier study, for consistency with a wider expanse of the nucleic acids literature we believed it was of value to compare the rates and yields of the reactions presented in Figures 1 when 100 mM NaCl is added to the reaction buffer, as buffers used in DNA experiments often contain 100 mM NaCl. The results of these higher salt concentration reactions are provided in the Supplementary Information (Figure S3). Briefly, the addition of 100 mM NaCl had minimal impact on the 9-mer substrate ligation kinetics. The initial rates, yields, and fits with double exponential curves are almost within experimental error of the results obtained without the additional NaCl. Similarly, the ligation reaction with the 5-mer substrate carried out at 4 °C exhibited negligible change in the reaction kinetics and overall yields. Significantly, reactions with the 5-mer substrate carried out at 25 °C and 37 °C exhibited an appreciable increase in ligation rates with added NaCl. The ligation rates at 25 °C increased to approximately those measured at 4 °C with or without NaCl, whereas even with the added NaCl the 37 °C reaction ligation rates remained low (i.e., <10% at the 24 hr time point). Given that the addition of 100 mM NaCl did not result in a substantial change in ligation rates or yields, the lower salt buffer was used for all other reactions reported in this study. Additionally, the lower salt buffer allowed easier analysis of the impact of the difference in stability of the 9-mer and 5-mer reaction systems on ligation rates and yields.

### Impact of substrate-to-hairpin ratios and side reactions on ligation efficiency

The reaction kinetics studies presented above all had a substrate:hairpin molar ratio of 10:1. This ratio was chosen based on preliminary studies which showed that substrate:hairpin molar ratios closer to 1:1 resulted in incomplete ligation at 24 hr for the 9-mer and the 5-mer substrates at all three reaction temperatures. Given the difference in stability between our 9-mer and 5-mer substrate-hairpin complexes, it was not obvious if these reactions did not go to completion because the ligation reaction kinetics were simply too slow with respect to the 24 hr time point, or because substrate oligonucleotides became a limiting reagent during the reactions, perhaps due to side reactions (e.g., cyclization) – a known problem with EDC activation chemistry (13,18).

Plots of ligation yields at the 24 hr reaction time point for various substrate:hairpin ratios are presented in Figure 3 for the 9-mer and 5-mer ligation systems. These plots show that reactions with the 9-mer system at 4 °C and 25 °C achieve maximum 24 hr yield (as indicated by the plateau) for substrate:hairpin ratios of 1.5:1 and greater (Figure 3). A higher ratio is required for the 37 °C reaction to reach the same level. For reactions with the 5-mer substrate, the 25 °C and 37 °C reactions do not reach their respective maximum possible 24 hr yields even at the 10:1 substrate:hairpin ratio. The 5-mer substrate reactions at 4 °C appear to be approaching a 24 hr yield plateau at the 10:1 ratio, but a significantly higher concentration is needed to actually reach maximum yield (see below). With the information provided by these plots, the reaction kinetic studies presented above, and additional analyses of side reactions (Supplementary Figure S4), we are able to gather considerable insight into why many reactions fall short of reaching complete ligation over the range of reaction conditions explored here and in other previously published studies.

**Figure 3.**
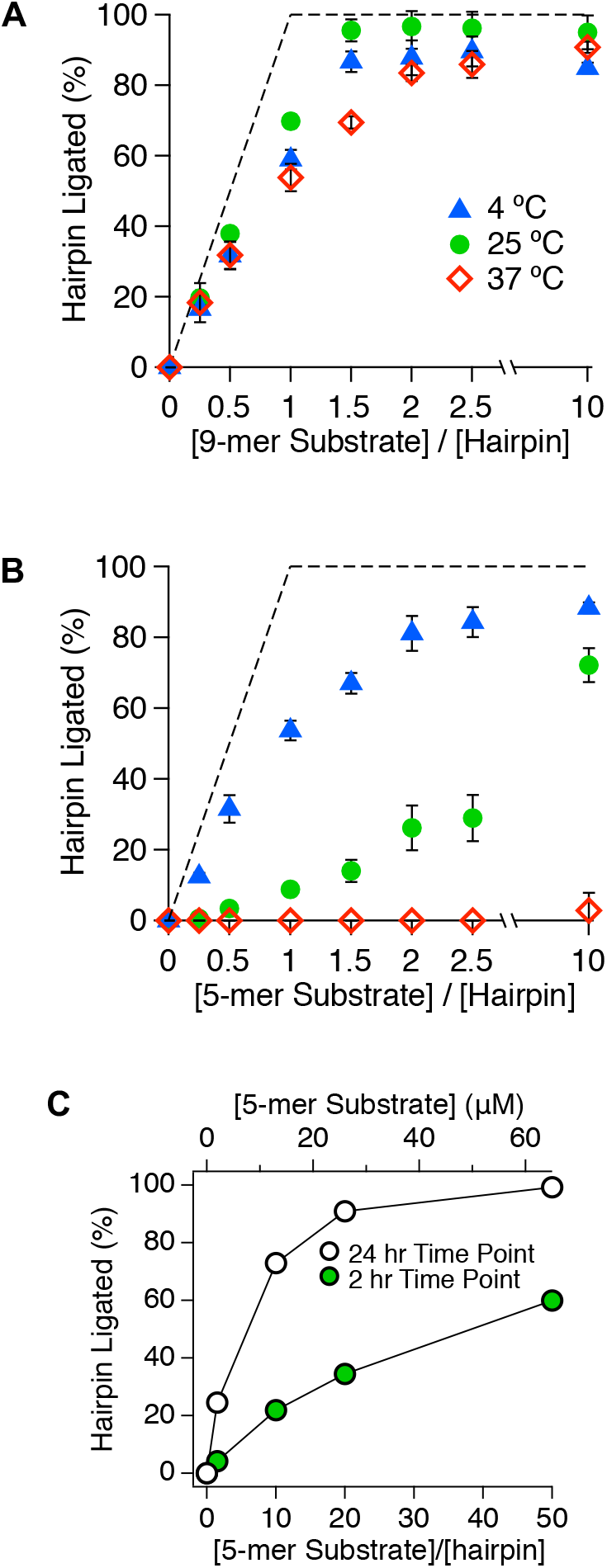
Effects of substrate-hairpin ratios on ligation efficiency for 3’-phosphate reactions. **(A)** Product formation for the 9-mer substrate as a function of relative molar concentrations of substrate to hairpin, after 24 hr reaction time. (**B**) Product formation for the 5-mer substrate as a function of relative molar concentrations of substrate to hairpin, after 24 hr reaction time. (**C**) Product formation for the 5-mer substrate comparing the 2 hr and 24 hr yields at 25 °C as a function of relative substrate:hairpin molar concentrations. Dashed lines in panels **A** and **B** indicate maximum possible yields for reactions as a function of substrate:hairpin ratios. All reactions contained 1.3 μM hairpin and were carried out in a buffer containing 5 mM MnCl_2_, 100 mM MES pH 6.0, and 250 mM EDC. Error bars represent the range of values measured for three separate experiments. Yields were obtained as described in the caption to Figure 1.

In the plots shown in Figure 3A and 3B the diagonal and horizontal dashed lines indicate where the data points would fall if the hairpins molecules in any of the experiments were quantitatively ligated to the substrate molecules within the 24 hr reaction time. The diagonal line corresponds to the region of the plot where the molar concentration of the substrate oligonucleotides is less than that of the hairpin oligonucleotides. The fact that the data points of all experiments fall below these dashed lines indicates that, for all the reactions studied, in no case are hairpins quantitatively ligated with substrates by 24 hr.

These variable substrate:hairpin ratio reactions, carried out at different temperatures, can be used to illustrate the three main factors that limit these reactions from producing their maximum possible ligation yields. The 4 °C data sets illustrate how the reaction temperature can make the chemical step of the reaction too slow for full ligation to occur even through full hairpin-substrate association is favored. As noted above, at 4 °C hairpins in the 9-mer reactions are fully associated with substrates, and there is also significant association for the 5-mer substrate reactions at 4 °C. However, the kinetics reaction plots for the 9-mer and 5-mer reactions run at 4 °C with substrate:hairpin ratios of 10:1 (Figure 1B and C) show that these reactions have not reached their maximum possible yields. Thus, the chemical steps of EDC-activated phosphodiester bond formation are simply too slow at 4 °C for the reactions to go to completion in 24 hr. This conclusion is also supported by 9-mer and 5-mer reactions with substrate:hairpin ratios of 1.25:1, also run at 4 °C, that show virtually the same reaction kinetics (Supplementary Figure S2).

The second reason that some reactions presented in Figure 3A and 3B do not reach their maximum possible yields, particularly those in the region where the substrate:hairpin ratio is less than one, is that substrate *and* hairpin oligonucleotides are likely being lost to side reactions with a rate that is comparable to the ligation rate. This possibility is perhaps best illustrated by the 9-mer substrate reactions run at 37 °C with a substrate:hairpin ratio of 10:1 which reach their plateau of maximum yield by 12 hr. These reactions fall short of complete hairpin ligation despite the rapid kinetics of these reactions and the reaction mixture containing 10-fold more substrate than hairpin (Figure 1B). An analysis of the 9-mer substrate concentration in an identical reaction as a function of time reveals that the concentration of unligated substrate decreases considerably more and faster than can be accounted for by ligation to the hairpin (Supplementary Figure S4). It is not clear how these substrates are being modified by EDC, but consistent with other reports (4,6,13), it appears that these “lost” or modified substrate are not able to ligate with the hairpin. This proposed contribution to the incomplete ligation of hairpins in the 9-mer substrate reactions at 37 °C is also supported by kinetic plots of reactions run with a substrate:hairpin ratio of 1.25:1, which have similar initial reaction rates as the 10:1 reactions, but are limited to a maximum yield of 65% regardless of reaction time (Supplementary Figure S2). We again emphasize that some fraction of hairpins are likely modified before ligation occurs, similar to the substrates modification, in a manner that precludes ligation, which would directly lower maximum possible ligation yields.

The third factor that limits full ligation by 24 hr is low equilibrium assembly of the hairpin-substrate complex. This factor appears to be the main reason that low yields are observed for the 5-mer substrate reactions run at 25 °C and 37 °C (Figure 3B). The low but near linear increase in the 24 hr yield of the 5-mer reactions run at 25 °C with increasing substrate concentration suggested to us that it might be possible to drive the hairpin to full assembly in particular ligation reaction, and the ligation rate to that of the chemical step, by increasing the substrate concentration to a value that is experimentally practical. In Figure 3C the 2 hr and 24 hr yields for this reaction are shown for substrate concentrations ranging from 1.25x to 50x relative to the 1.3 μM hairpin concentration. Importantly, the 2 hr yield of the 5-mer reaction with 50x substrate is 60%, which is very close to the 63% 2 hr yield observed for the 9-mer reaction when also run 25 °C. Thus, the 5-mer reactions run at 25 °C with 50x substrate reach a ligation rate consistent with fully assembled hairpins with substrates. Also consistent with this conclusion, note that the 24 hr ligation yield for the 50:1 5-mer substrate:hairpin reaction is 99% (Figure 3C), which is slightly higher than that observed for the 9-mer ligation reactions run at 25 °C for a substrate:hairpin ratio of 1.5:1 to 10:1 (Figure 3A), which are all expected to have fully assembled substrate-hairpin complexes.

Based on the results presented thus far, we can conclude that several convoluting factors are governing the reaction yield, including equilibrium assembly of the substrate:hairpin complex, rate of the chemical step, side reactions, and half-life of EDC (4,14), all of which will have a temperature dependence.

### A model for determining concentrations required for high ligation yields

Using information gained from our ligation systems we attempted to construct a mathematical model to predict the yield that would be obtained for a particular experimental system at any given hairpin concentration and substrate:hairpin ratio. The model developed requires only three parameters, the substrate-hairpin equilibrium constant (*K*_2_, Scheme 1C), the kinetic rate constant for ligation bond formation (*k*_2_, Scheme 1D), and the kinetic rate constant of substrate loss (*k*_4_, Scheme 1F). Full derivation of this model is provided in Supplementary Information.

In Figure 4 we show a contour plot of the predicted ligation yields for the 5-mer 3’-phosphate substrate reaction run at 25 °C for 24 hr as a function of *K*_2_ and hairpin concentration (their product being the independent variable of the horizontal axis) versus the ratio of substrate and template concentrations (this ratio being the independent variable of the vertical axis). This plot is particular to the 5-mer ligation reaction at 25 °C; the kinetic rate constant for bond formation (*k*_2_ = 0.49 hr^−1^) was determined from 9-mer ligation reactions carried out at 25 °C, and the kinetic rate constant for substrate loss (*k*_4_ = 0.103 hr^−1^) is based on the experimentally determined kinetics of the 5-mer ligation system. Briefly, the values for *k*_2_ and *k*_4_ were determined by curve fitting of the kinetic ligation data shown in Figure 1B and 1C for reactions with a 10:1 substrate:hairpin ratio. The RMSD best fits of this data also revealed that *K*_2_ of the 5-mer reaction at 25 °C is 2.8 ×10^4^ M^−1^ (see Supplementary Information for details of parameter determination). Because these values are based on experimental data, a single point on the contour plot of Figure 4 is an experimentally determined value upon which all the yields of all other points are determined using our model. This single data point is shown as a green filled circle with a horizontal coordinate of 0.036, (*K*_2_ x [hairpin] = 2.8 x10^4^ M^−1^ × 1.3 μM), and a vertical coordinate of 10 ([substrate]/[hairpin] = 13 μM/1.3 μM).

**Figure 4.**
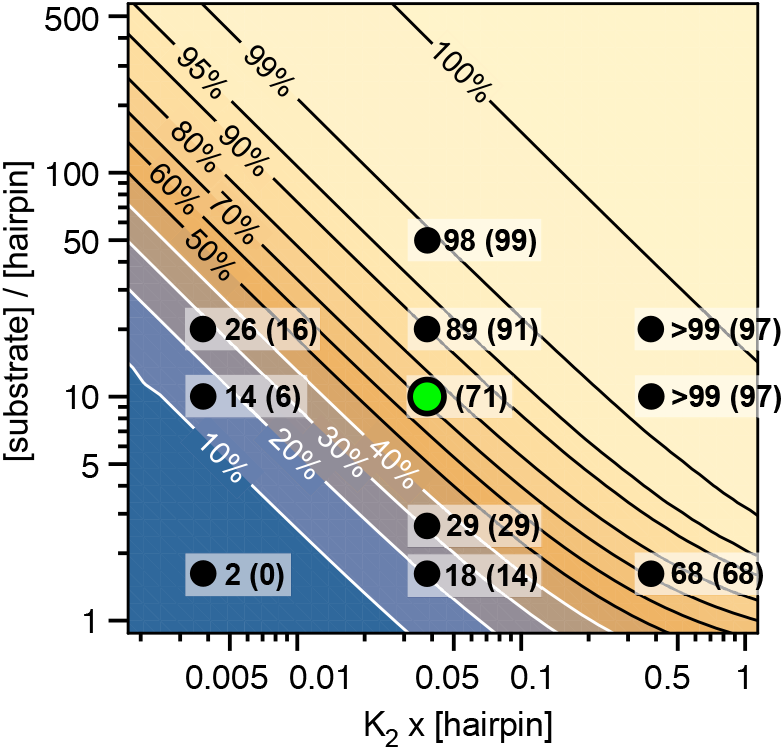
Contour plot showing predicted and experimentally determined ligation yields for the 5-mer 3’-phosphate substrate at 25 °C after 24 hr in terms of the dimensionless independent variables equilibrium constant (*K*_2_) x[hairpin] and [substrate]/[hairpin]. The values outside the parentheses are model-predicted yields; values inside parentheses represent experimental yields. The single green point corresponds to the experimental system from which the equilibrium constant and the substrate loss rates were determined.

Figure 4 was formulated to predict the ligation yield for the 5-mer system run at 25 °C at the 24 hr for any substrate and hairpin concentration. To test the accuracy of our model we placed points on the contour plot from other 5-mer ligation experiments presented above. For the same horizontal value of *K*_2_ x [hairpin] = 0.036, we have plotted points corresponding to 24 hr 5-mer ligation experiments in which substrate:hairpin ratios of 1.5, 2.5:1, 20:1 and 50:1 were used (all run at 25 °C, with [hairpin] = 1.3 μM). The predicted 24 hr yield for each of these experiments are provided next to their corresponding point on the contour plot, along with the experimentally determined yields in parentheses. All four of the experimental yields with *K*_2_ x [hairpin] = 0.036, ranging from 14% to 99%, are within error of their predicted values.

Experiments were also carried out to test the accuracy of our model at different locations along the horizontal axis. Because the plot in Figure 4 is particular to the reaction conditions of the data set upon which the model parameters were derived, it would be difficult to change the equilibrium constant, *K*_2_, without the possibility of also changing other model parameters. For example, *K*_2_ can be changed by increasing or decreasing temperature, but a change in temperature would also alter the rate of the chemical steps of ligation. Thus, we changed the horizontal positions of our experiments by increasing and decreasing the hairpin concentration from the original data point (green dot). For these experiments 24 hr ligation yields were measure for hairpin concentrations of 0.13 μM and 13 μM with the 5-mer substrate:hairpin ratios of 1:1, 10:1, and 20:1. At high hairpin concentrations (*K*_2_ x [hairpin] = 0.36) the model predicted yield that were virtually identical to experimental yields. At low hairpin concentrations (*K*_2_ x [hairpin] = 0.00364) the experimental yields were somewhat lower than predicted yields. It is possible that the predicted lower yields differ more from their corresponding experimental values because experimental errors can be greater when measuring low amounts of product formed, or that the model does not sufficiently capture the impact of substrate and hairpin modifications in low concentration reactions where most of the hairpins are not associated with substrates. Nevertheless, the fact that all the experimental and predicted yields in Figure 4 are within experimental error provides support for our rather simple model being able to predict ligation yields over a wide range of hairpin and substrate concentrations, for a particular system.

To apply the model developed to construct the contour plot in Figure 4 to other ligation systems it will be necessary to determine the kinetic rate constant for bond formation, *k*_2_, and the kinetic rate constant for substrate loss, *k*_4_, for each ligation system. It is possible that the kinetic rate constant for bond formation may be the same for stable complexes using the same activation chemistry, as appears to be the case for the 9-mer and 5-mer systems studied here. However, the kinetic rate constant for substrate loss should be expected to vary tremendously. Richert and co-workers showed that the loss of dimer and trimer substrates to cyclization could be the main inhibitor of chemical ligation (13). Even in our system, where substrate cyclization does not appear to be severely inhibiting the ligation reaction, we still detect a loss of substrate possibly due to other modifications by EDC, with the 9-mer substrate showing greater rates of modification than the 5-mer substrate (Supplementary Figure S4). Other factors discussed above, such as phosphate position, substrate-hairpin stability enhancement by EDC, organic catalysts in reaction buffer, and nucleotide sequence (discussed below) are also expected to impact the kinetic rate constants and ligation-active complex stability of a particular system.

### Influence of nick-site flanking base pairs on ligation initial rates and 24 hr yields

Thus far we have utilized the single hairpin sequence and its complementary substrates shown in Figure 1, which has the nick site flanked by two cytosine bases. The specific base pairs flanking a nick site have been previously observed to impact ligation yields (7,8,13). It was therefore important to also use our test system to explore the sensitivity of EDC-activated phosphodiester bond formation to the identity of base pairs flanking a nick site, having demonstrated that we have a ligation test system with stable pre-ligation substrate-hairpin complexes. For these studies we purchased an additional seven hairpins and their complementary 9-mer substrates to create nick sites with the bases being four possible GC base steps (3’-C pC −5’; 3’-C pG −5’; 3’-G pC −5’ and 3’-G pG −5’) and the four possible AT base steps (3’-T pT −5’; 3’-T pA −5’; 3’-A pT −5’ and 3’-A pT −5’). The 9-mer substrate was chosen at a substrate:hairpin ratio of 1.5:1 to ensure that maximum ligation could be obtained after 24 hrs at most temperatures, as illustrated by the data presented in Figure 3A. The substrate-hairpin sequences with AT base pairs flanking the nick site were adjusted so that they would have the same %GC content, and therefore similar stability, as the set with GC base pairs flanking the nick site, although it was appreciated that the base pair stability and stacking interactions at the nick site could vary by sequence.

As shown by the curves in Figure 1B, yields measured at the 2 hr time point are a good balance between sufficient product for accurate quantitation of the slowest 9-mer ligation (at 4 °C), while still providing sufficient dynamic range to discern differences in the faster rates, such as between 9-mer ligation at 37 °C as compared to 25 °C. Thus, for this set of ligation experiments we used the yields measured at the 2 hr reaction time point as a semi-quantitative means for comparing the initial rates of different sequence and the yields measured at the 24 hr time point as an estimate of long reaction time yield. The comparison of yields for these two reaction time points for the four possible GC base pair steps at the nick site reveals a striking difference between the ligation rates of the C pC and G pG flanking base pairs (Figure 5A). For example, the initial rate of ligation at a C pC nick site is three times the initial rates of a G pG nick site at 4 °C and 25 °C. For reaction carried out at 37 °C this difference in the initial rates is decreased but remains significant. Because all substrate:hairpin combinations had similar predicted T_m_’s, we expect that the differences observed in initial ligation rates is due to differences in base stacking near the activated phosphate and the alcohol nucleophile. We hypothesize that presence of the GG stacked base pairs induces rigidity on the bases and slows ligation by holding the reactive groups away from their optimal reaction geometry, particularly at lower reaction temperatures. This rigidity can be overcome by increasing the temperature of the reaction, which allows the oligonucleotides access to more conformations. Nevertheless, the maximum yield attained by the G pG system is consistently lower than those of the other system, even at 37 °C.

**Figure 5.**
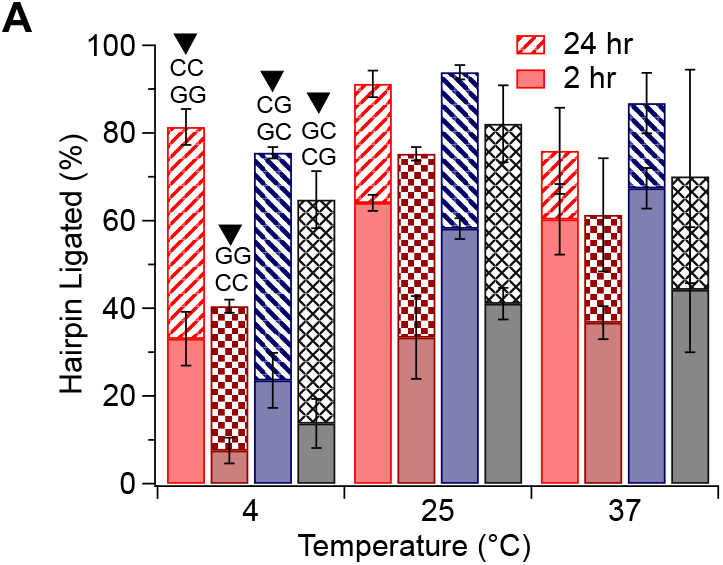
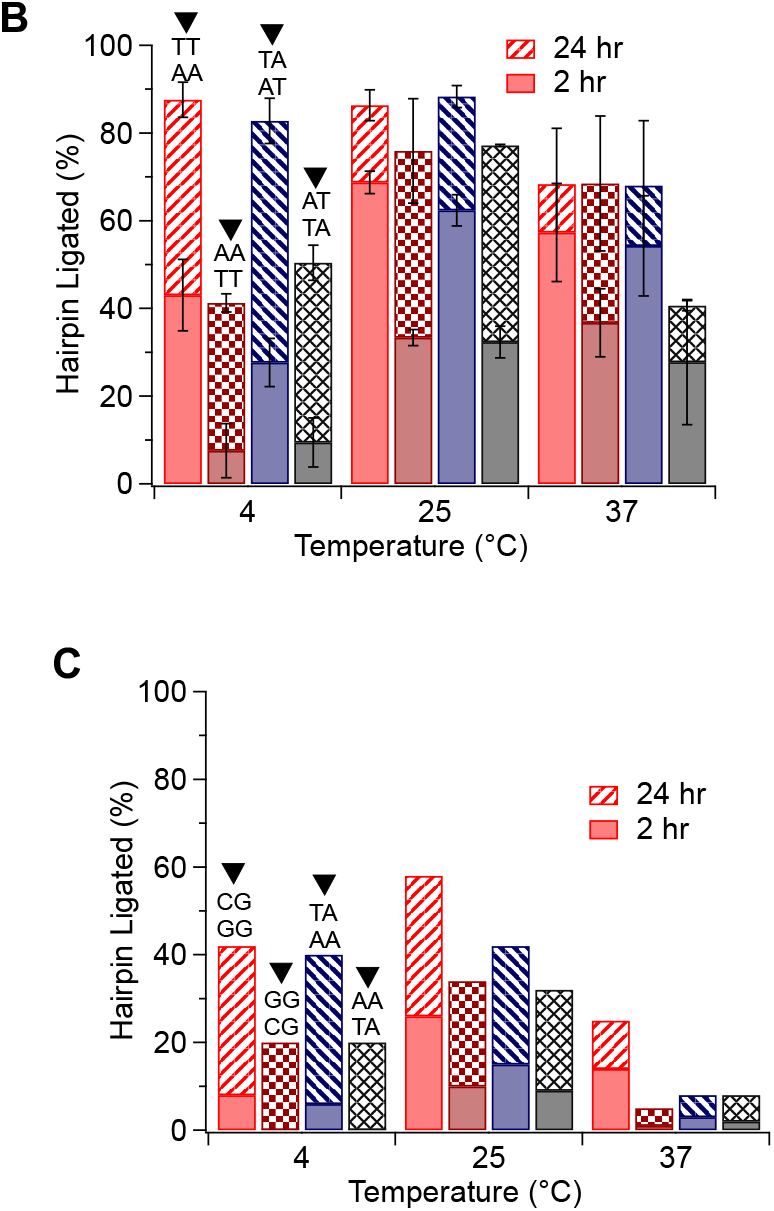
Effects of nick-site flanking base pairs on ligation efficiency **(A)** Watson-Crick base pairs product formation for the CG substrate:hairpin system at different temperatures. **(B)** Watson-Crick base pairs product formation for the AT substrate:hairpin system at different temperatures. **(C)** Mismatch base pairs product formation for both the CG and AT substrate:hairpin system at different temperatures. All reactions were carried out for 2 hr and 24 hr indicated by the unfilled and shaded bars respectively, at 4 °C, 25 °C, and 37 °C, with a substrate:hairpin ratio of 1.5:1 in a buffer containing 5 mM MnCl_2_, 100 mM MES pH 6.0, and 250 mM EDC. Directionality of sequences is shown in Scheme 1H. Each bar graph is from an average of two or three independent experiment replicates; with error bars representing the range of values measured. Data points were obtained as described in Figure 1 caption.

Similar results are observed for the four possible AT base steps that flank the nick site (Figure 5B). The direct correspondence between GC and AT systems for the impact of pyrimidine-pyrimidine stacking at the flanking site, versus purine-purine stacking, support the hypothesis that these differential initial rates and temperature dependence is due to base stacking, as opposed to non-Watson-Crick hydrogen bonding interactions that would be expected to be different for the GC and AT base steps. These findings are similar to those observed by Carrierio, and Damha (9) in which the G pT ligation site gave 10% yield, while T pT site gave 66% yield, and others (20,21), that have also found that pyrimidine-pyrimidine base flanking is better than other base flanking compositions.

We also decided to examine if purine-purine and pyrimidine-pyrimidine stacking around a nick site would have a similar impact on ligation at mismatched base pairs. In Figure 5C it is shown that when a purine-pyrimidine flanking base pair is used, such that the purine stacking is opposite the substrate:hairpin nick, 40% of ligated products are observed at 4 °C. In contrast, when a purine-purine flanking base pair is used with a purine mismatch, the yield is decreased by half to 20% at the same temperature. As observed previously, an increase in reaction temperature to 25 °C led to an increase in initial rates and maximum yields. However, at 37 °C, the incorporation of mismatches is significantly reduced to less than 10% in most cases. This drastic decrease in reactivity at 37 °C can be attributed to the lower melting temperature the mismatch nucleotide introduces to the system (i.e. the T_m_ of the main substrate:hairpin sequence decreases from 45 °C to 37 °C) (Figure S5). It is interesting to note that the incorporation of mismatched purines with the purine-pyrimidine flanking base pair (Figure 5C) is identical in initial rates and maximum yield to the incorporation of Watson-Crick purine-purine flanking base pairs (Figures 5A and B). These findings also corroborate studies which have previously found that chemical ligation is tolerant to the introduction of mismatches (7,22).

## CONCLUSION

Using a well-defined bimolecular system we have explored how ligation rates and yields using the carbodiimide EDC as an activating agent are impacted by several experimental parameters. Most fundamentally, our results demonstrate quantitatively how the equilibrium association constant of a substrate:template complex directly impacts the initial ligation rate and overall yield (i.e., long incubation time yield). The substantial temperature dependence of the chemical step(s) of phosphodiester bond formation by EDC activation presents a challenge for selecting the optimal temperature for carrying out chemical ligation. In particular, if a substrate:template complex has significant stability, then increasing the temperature of the ligation reaction will allow reaction kinetics and, in some cases, higher yields. However, if a substrate:template complex has low stability, then lowering the reaction temperature may be necessary to reach maximum ligation yield at the expense of lower rates of bond formation. This conclusion may seem obvious, and previous investigators have similarly concluded that the efficiency of chemical ligation is dependent on the efficiency of hybridization and ligation rate (8). However, our goal in this work has been to provide a quantitative framework for optimizing ligation yields. It is not always clear *a priori* what is the optimal temperature for chemical ligation, even with theoretical prediction or experimental measurements of assembly stability, as the presence of a condensing agent and substrate activation can alter association constants. Additionally, relatively small changes in temperature, such as 10 °C, can move a system from near optimal ligation yields to very low yields. In the very least, the consideration of these parameters and measuring the kinetics of representative reactions can be sufficient to determine the conditions necessary to achieve near optimal yields within a reasonable time frame.

As has recently been shown to be the case for reactions involving the template-directed ligation of dimers and trimers (13), our work with longer substrate oligonucleotides has also shown that the substrates and template modification by EDC can prevent a reaction from going to completion. In addition to ligation yields, it is expected that for many applications, DNA modification will be an unwanted outcome of a ligation reaction. Thus, in addition to optimizing reactions to reach higher yields, having a ligation reaction come to completion within the shortest amount of time may be the best way to minimize unwanted modifications of the target products.

For application where there is some flexibility in DNA sequence and terminal phosphate position, the studies presented here provide clear guidance for how to design nick sites that will be most amenable to chemical ligation. Our results have shown that having the terminal phosphate on the 3’ end of one of the two oligonucleotides to be ligated is far more productive than having the phosphate on the 5’ end of the other oligonucleotide. The substantial difference we observe in the rates of bond formation can result in a difference in maximum yield being 95%, in the case of a 3’ phosphate, versus 40% for precisely the same DNA system, except for the placement of the phosphate on the 5’ terminus of the nick site. It is important to note that the opposite appears to be true for RNA ligation (6,14,23).

The model reactions presented here have also revealed the substantial impact that the identity of nick site flanking base pairs can have on ligation rates and yields. The general observation that a pyrimidine-pyrimidine step at a nick site, either C C or T T, leads to higher ligation rates than a purine-purine step, either G G or A A, suggests that the energetics of purine stacking can place the reactive groups as a nick site in relative position or orientations that are more or less favorable for bond formation. This local impact of sequence on ligation yields can be minimized by increasing the reaction temperature, but this does present another temperature-dependent parameter that must be considered in optimizing a ligation reaction, along with substrate-template association constant and the kinetic of the chemical step.

With respect to using chemical ligation in systems that contain more than one pair of oligonucleotides to be ligated, such as in the creation of covalent DNA nanostructures and in the construction of model self-replicating systems of increasing complexity, the data presented here raises a caution. Like early work by James and Ellington (7), our results have shown that chemical ligation is not nearly as selective for Watson-Crick base pairs as is the case with ligase enzymes. Nonetheless, chemical ligation offers several advantages over enzymatic ligation including the ability to ligate non-natural base pairs. The promiscuity of chemical ligation of assemblies containing mismatched bases is necessary for the creation of non-regular DNA structures, but it could also result in unplanned cross-reaction of substrates as well as the introduction of mismatched base pairs in products. Operating a ligation reaction at a higher temperatures can potentially make the system more selective for Watson-Crick base pairing, but again the sensitivity of a reaction to increasing temperature must be considered, as it may not be possible to operate at a temperature that minimizes undesired cross-reactions while still achieving high yields for the desired products.

Finally, we note that the parameters and models provided in this work for optimization of DNA chemical ligation are not limited to a bimolecular system, such as the test system we used here, but can also be applied to any template-directed ligation system for which equilibrium constants of substrate association to form a ligation-active complex can be determined.

## SUPPLEMENTARY DATA

Supplementary Data are available at NAR online.

## ACKNOWLEDGEMENT

The authors would like to acknowledge A.L. Colinas, Dr. D.M. Fialho, and Prof. Roger Wartell for helpful discussions. We thank Dr. Bradley Burcar for technical assistance, and the Petit Institute for Bioengineering and Biosciences for use of core facilities.

