## Supplementary Information for "Impact of substrate-template stability, temperature, phosphate location, and nick-site base pairs on non-enzymatic DNA ligation: Defining parameters for optimization of ligation rates and yields with carbodiimide activation"

This file includes:

Supplementary Materials and Methods
DNA sequences
Supplementary Figures S1-S6
Supplementary Table S1
Details on Model Construction
Supplementary References
Model Codes in Matlab and Mathematica (in separate files)

### Supplementary Materials and Methods

Melting temperature ($T_m$) determination

UV absorbance was used to monitor the thermal denaturation of the hairpin and complementary short oligonucleotide complex. DNA samples were prepared with 5 mM $MnCl_2$, and 100 mM MES pH =6.0, with a template hairpin concentration of 1.3 μM, and a substrate concentration of 2 μM. UV measurements were performed on 10 mm quartz cuvettes in a temperature-controlled UV-Vis spectrophotometer (Cary Agilent UV-Vis Multicell Peltier) with nitrogen flowing through the sample chamber at low temperatures. To determine $T_m$ values, heating and cooling traces were generated for each sample by recording spectra at 260nm from 18 to 58 $^oC$ at intervals of 1 $^oC$. $T_m$ values were determined as described by Mergny & Lacroix (1).

Side product formation reactions

For monitoring the formation of side products, the ligation experiments were carried out as described previously in the main text. At the indicated time point, a 10 μL volume of the reaction mix was quenched with an equal amount of formamide (Alfa Aesar, 99.5%) and stored at -80 $^oC$. The decay in the substrate oligonucleotides was then monitored using both HPLC and mass spectrometry as described below.

HPLC Analysis

All HPLC analysis was conducted on an Agilent 1260 Infinity HPLC system. A DNA PAC$^{TM}$ PA 200 4 μm Anion Exchange Column (4x250 mm) was used for analysis. The following buffers and conditions were used for all analyses: (A) 12.5 mM Tris, pH 8.0; (B) 12.5 mM Tris, pH 8.0, 1.5M NaCl. Flow rate 1.20 mL/min, 10 μL sample injection, and column at 60 $^oC$. Absorbance was measured at 260 nm and calculation of peak areas were carried out using Chemstation B.04.03.

The following elution scheme was used: 0-5 minutes used isocratic flow at 95% (A), 5.1-20 minutes used gradients from 85-55% (A), 20-28 minutes used gradients 55-20% (A), and 95% (A) from 28.1 – 31 minutes. Peak assignments were conducted by running oligonucleotide standards and confirming new peaks through spiking and MS analysis.

**DNA Sequences**

Hairpin template

(Main) CG Hairpin: 5´ CAGTCACGGAACGTGACTGGACAGGAGA 3´ 6-FAM
GC Hairpin: 5´ GAGTCACGGAACGTGACTCCACAGGAGA 3´ 6-FAM
GG Hairpin: 5´ GAGTCACGGAACGTGACTCGACAGGAGA 3´ 6-FAM
CC Hairpin: 5´ CAGTCACGGAACGTGACTGCACAGGAGA 3´ 6-FAM
AT Hairpin: 5´ACGTCACGGAACGTGACGTTACAGGAGA 3´ 6-FAM
TA Hairpin: 5´TCGTCACGGAACGTGACGAAACAGGAGA 3´ 6-FAM
TT Hairpin: 5´TCGTCACGGAACGTGACGATACAGGAGA 3´ 6-FAM
AA Hairpin: 5´ACGTCACGGAACGTGACGTAACAGGAGA 3´ 6-FAM

Substrates

(Main) 9-mer: 5´ TCT CCT GTCp 3´
(Main) 5-mer: 5´ CT GTCp 3´

Other substrates

G 9-mer: 5´ TCT CCT GTGp 3´
A 9-mer: 5´ TCT CCT GTAp 3´
T 9-mer: 5´ TCT CCT GTTp 3´

The sequences highlighted in green are in the loop of the hairpin.

### Supplemental Figures

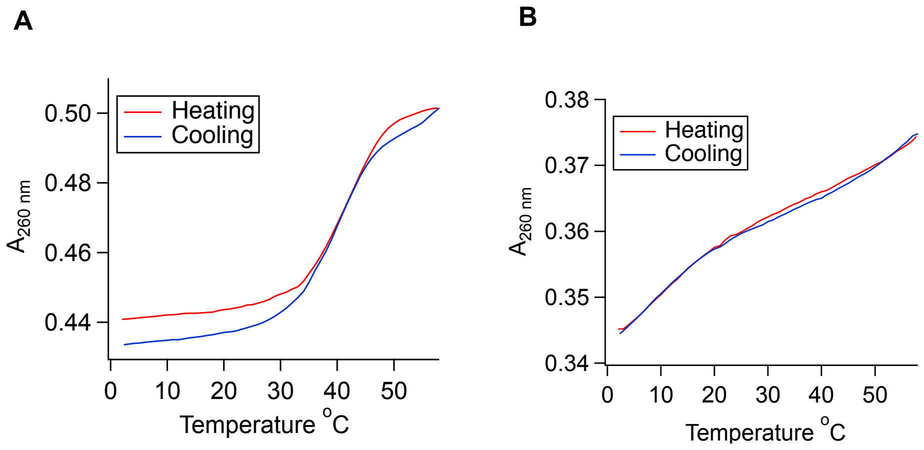


**Figure S1**: Determination of melting temperature ($T_m$) of the substrate-hairpin assembly by monitoring absorption at 260 nm as a function of temperature. (**A**) The 9-mer dissociates from the hairpin at a $T_m$ of 45 °C. (**B**) No clear transition of the 5-mer hairpin dissociation can be observed over the temperature range of 2 °C and above. However, the change in slope observed at 20 °C is consistent with a $T_m$ around the predicted value of 6 °C. Note that hairpin secondary structure has a $T_m$ of approximately 85 °C.

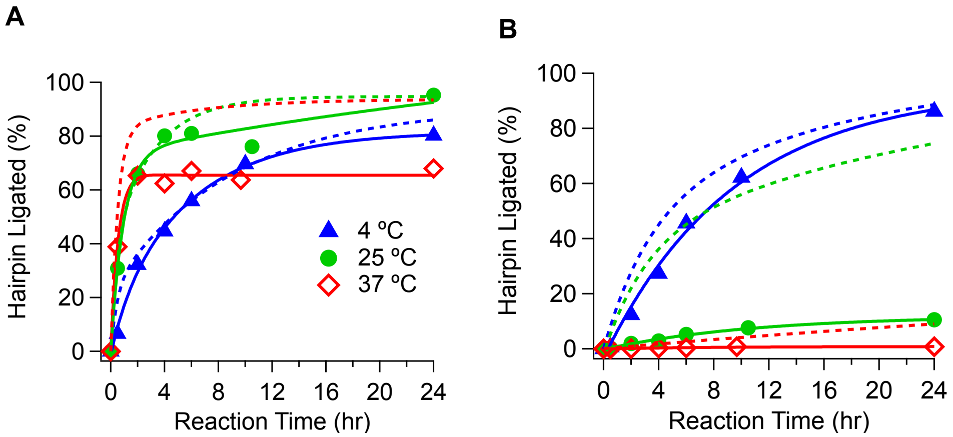


**Figure S2**: Kinetics of chemical ligation with a substrate:hairpin ratio of 1.25:1. (**A**) Kinetic data for 3'-phosphate 9-mer substrate. (**B**) Kinetic data for the 3'-phosphate 5-mer substrate. For all experiments hairpin concentration was 1.3 μM. Buffer conditions: 5 mM $MnCl_2$, 100 mM MES, pH 6.0, and 250 mM EDC. Markers are experimental data, solid lines are double exponential fit of data to guide the eye. Dashed lines are double exponential fits of data when reaction was conducted using a substrate:hairpin ratio of 10:1 (data shown in Figure 1, main text).

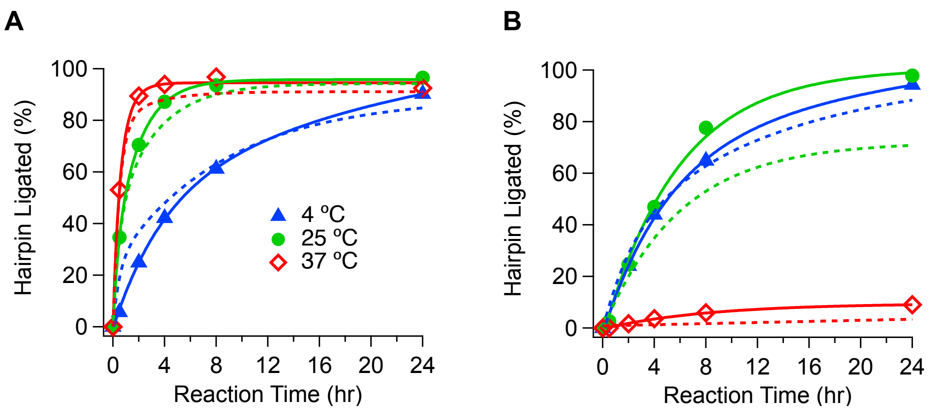


**Figure S3**: Impact of adding 100 mM NaCl to the ligation reaction buffer. (**A**) Kinetic data for hairpin ligation with the 3'-phosphate 9-mer substrate. (**B**) Kinetic data for hairpin ligation with the 3'-phosphate 5-mer substrate. Markers and solid lines are yields observed with 100 mM NaCl added to the ligation buffer. Dashed lines are double exponential fits of data for corresponding reactions in standard reaction buffer without the addition of NaCl. Standard reaction buffer: 5 mM $MnCl_2$, 100 mM NaCl, 100 mM MES, pH 6.0, and 250 mM EDC. Reaction temperatures as indicated in **A**.

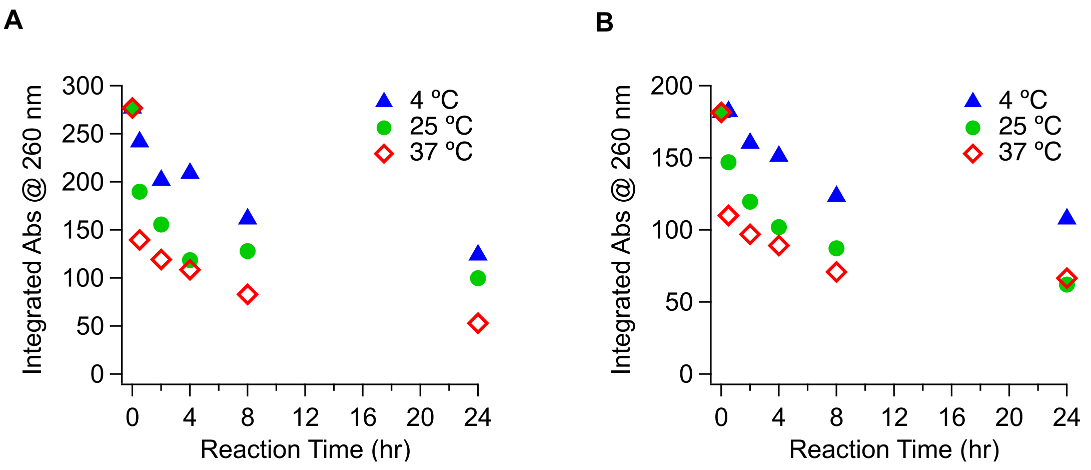


**Figure S4**: Kinetics of substrate decay in ligation reaction buffer using integrated absorbance from UV-HPLC. (**A**) Kinetic data for 3'-phosphate 9-mer substrate. (**B**) Kinetic data for the 3'-phosphate 5-mer substrate. All reactions had a 10:1 substrate:hairpin molar ratio, and 1.3 μM hairpin concentration. Buffer was 5 mM $MnCl_2$, 100 mM MES, pH 6.0, and 250 mM EDC. Reaction temperatures as indicated in graphs.

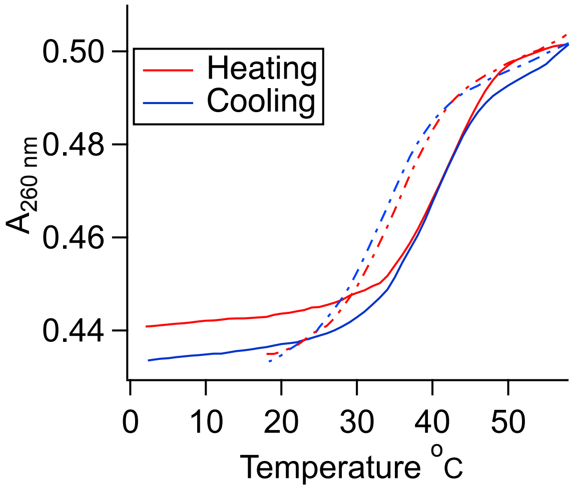


**Figure S5**: Melting temperature ($T_m$) determination of the substrate-hairpin association. The dashed lines represent the mismatch 9-mer substrate, while the solid lines represent the $T_m$ of the Watson-Crick 9-mer substrate shown in Figure S1. The mismatch 9-mer dissociates from the hairpin at a $T_m$ of 37 °C.
.

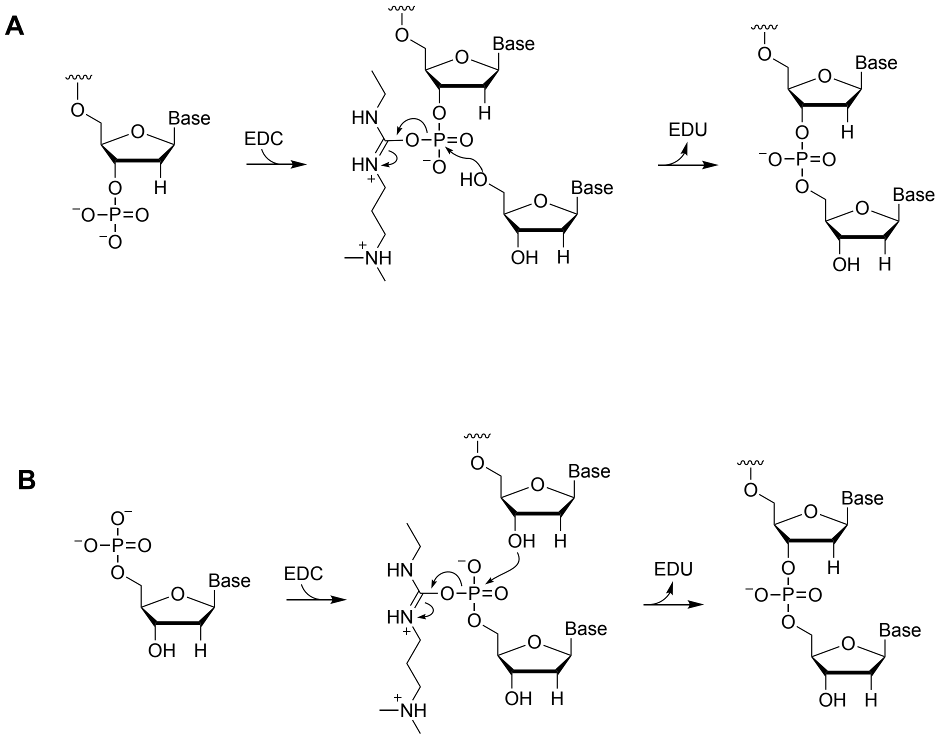


**Figure S6**: Key steps in the reaction mechanism for substrate activation by EDC and subsequent ligation. (**A**) Reaction pathway for the 3´p reaction (**B**) Reaction pathway for the 5´p reaction.

**Supplemental Table**

**Table S1.** Predicted fraction of hybridized substrates at different reaction temperatures for a solution containing a substrate:hairpin ratio of 10:1 and 1.5:1. These values are predicted for a solution containing 5mM $MgCl_2$ and 100mM Tris buffer.

| | 5-mer | | 9-mer | |
|---|---|---|---|---|
| Excess substrate | 1.5 x | 10x | 1.5 x | 10x |
| Reaction Temperature (°C) | Hybridized substrate ($x$) | Hybridized substrate ($x$) | Hybridized substrate ($x$) | Hybridized substrate ($x$) |
| 4 | 0.18 | 0.6 | 1 | 1 |
| 25 | $4.0 \times 10^{-3}$ | 0.023 | 0.94 | 0.99 |
| 37 | $4.6 \times 10^{-4}$ | $2.8 \times 10^{-3}$ | 0.36 | 0.86 |

**Details on Model Construction**

1. <u>Estimating the fraction of bound substrates at different reaction temperatures</u>

Since the $T_m$ of the 5-mer could not be measured, the "MELTING" program (2) based on work by Santa Lucia et. al (3) was employed. The predicted $T_m$ of the 5-mer was -4 $^oC$ and 6 $^oC$ at an excess concentration of 1.5x and 10x respectively, while the predicted $T_m$ of the 9mer was 35 $^oC$ and 45 $^oC$ at an excess concentration of 1.5x and 10x respectively. The conditions between the experimental melts and the predicted melts were slightly different since the buffer for the predicted melts was assumed to be 5 mM $MgCl_2$, Tris pH = 7.0, which is different from the ligation buffer conditions of 5 mM $MnCl_2$, MES pH = 6.0 in which Figure S1 was calculated. Therefore, the discrepancy between the predicted and experimental $T_m$ of the 9-mer can be expected. Nonetheless, since the experimentally observed melts for the 9-mer is above the predicted melts from the "MELTING" program, the number of bound substrates will more likely be under predicted rather than over predicted.

Using the predicted enthalpy from the "MELTING" program (i.e. - 261,668 J/mol for the 9-mer, and -136,686 J/mol for the 5-mer), the ratio between the equilibrium constant at the melting temperature was found at the different reaction temperatures according to Van't Hoff Equation S1.

$$\ln \frac{K_{eq,Tm}}{K_{eq,T}} = \frac{\Delta H^o}{R} \left(\frac{1}{T} - \frac{1}{Tm}\right) \quad \text{(S1)}$$

The value of $\frac{K_{eq,Tm}}{K_{eq,T}}$ was then used to find the fraction bound at different reaction temperatures, for different excess amounts according to Equations S2 & S3.

$$K_{conc,T} = \frac{x}{(A-x)(B-x)} \quad \text{(S2)}$$

where $x$ = concentration of hybridized substrate, = 0.5 at $T_m$
$A$ = Initial hairpin concentration
$B$ = Initial substrate concentration

$$\frac{K_{eq,Tm}}{K_{eq,T}}(pred) = \frac{K_{conc,\,Tm}}{\frac{x_T}{(A-x_T)(B-x_T)}} \quad \text{(S3)}$$

$K_{conc,T}$ is found at the $T_m$ and used to predict the fraction of hybridized substrate $x_T$ at different temperatures, according to Equation S3. These results are summarized in Table S1.

2. <u>Modeling of the ligation reaction</u>

The rate of ligation was modeled using second order kinetics shown in the reaction scheme below.

$$\begin{array}{l} H + O \underset{k_f/K_{eq}}{\overset{k_f}{\rightleftharpoons}} I \xrightarrow{k_c} P \\ \quad\;\; {\scriptstyle k_S}\downarrow \\ \quad\;\;\; S \end{array}$$

**Scheme 1**. Reaction network of ligation reaction.

where H represents the concentration hairpin, O is the complementary oligonucleotide, I is the hybridized intermediate complex, P is the ligated products, and S is the oligonucleotides side products. The four rate constants are forward hybridization rate constant $k_f$, backward hybridization rate constants, equilibrium constant $K_{eq}$, side product formation rate constant $k_s$, and chemical ligation rate constant $k_c$. The side reactions were included in the model based on our experimental results as well as past reports (4,5).

The overall ligation reaction scheme is defined by the differential equations:

$$\frac{dS}{dt} = k_s O \qquad \text{(S4)}$$

$$\frac{dI}{dt} = k_f(HO - \frac{I}{K_{eq}}) - k_c I \qquad \text{(S5)}$$

$$\frac{dP}{dt} = k_c I \qquad \text{(S6)}$$

$$\frac{dH}{dt} = -k_f(HO - \frac{I}{K_{eq}}) \qquad \text{(S7)}$$

$$\frac{dO}{dt} = -k_f(HO - \frac{I}{K_{eq}}) - k_s O \qquad \text{(S8)}$$

For the contour plot generated in the main text, the rate constants in Equations S4-S8 are related as follows, $k_c = k_2$, $k_s = k_4$, and $K_{eq} = K_2$. The following steps were taken to obtain the rate constants;

1. The value of the chemical ligation rate constant $k_c$ for the 5-mer reaction was obtained by fitting the 9-mer experimental kinetic data (Figure 2B main text, 25 °C) with Equations S4-S8. $k_f$ was set to $1*10^{10}$ $hr^{-1}M^{-1}$ to ensure the model depended only on the equilibrium constant which was found for the 9-mer using Equation S1. During this optimization, the side reaction rate constant for the 9-mer reaction was also obtained. The model revealed that $k_c = 0.4856$ $hr^{-1}$and this value was assumed to be equal for both the 5-mer and 9-mer reactions as discussed in the main text. The Matlab and Mathematica files to solve these equations are titled "Ligationconstant_solver" and "Ligationconstant" respectively.

2. Next, the equilibrium constant $K_{eq}$ and the side reaction rate constant $k_s$ for the 5-mer reaction were found for the optimized chemical ligation rate constant ($k_c = 0.4856$ $hr^{-1}$) by fitting Equations S4-S8 with experimental values obtained in Figure 2C main text. The values of these constants are as follows, $K_{eq} = 2.82*10^4$ $M^{-1}$, and $k_s = 0.103$ $hr^{-1}$. $k_f$ was kept constant at $1*10^{10}$ $hr^{-1}M^{-1}$, similar to step 1. The Matlab and Mathematica files to solve these equations are titled "Equilibriumconstant_solver" and "Equilibriumconstant" respectively.

3. After the rate constants were determined for the 5-mer reaction system (Figure 2C main text, 25 °C), a general plot was made with varying substrate:hairpin ratios and equilibrium constants. Experiments were then carried out at different substrate:hairpin ratios and equilibrium constants and compared to the predicted value on the contour plot. The Matlab and Mathematica files to make the contour plot are both titled "Contourplot".
